# Molecular methods reveal responses of bacterial communities, including indicator species, to ballast water management

**DOI:** 10.64898/2026.04.23.720405

**Authors:** Sarah A. Brown, Scott P. Keely, Nichole E. Brinkman, Olivier Laroche, Anastasija Zaiko, Vanessa Molina, Matthew First, Lisa Drake, John A. Darling

## Abstract

Ballast water (BW) is an important vector for the global translocation of bacterial taxa, including pathogens. Legal frameworks establishing limits on the discharge of live organisms into recipient environments have been designed to reduce risks of microbial invasions, and understanding the impacts of BW management on bacterial communities is critical for assessing the effectiveness of these practices. Here we evaluate changes in bacterial communities associated with both BW treatment (BWT) and a combined management approach of BWT plus BW exchange (BWT+E). Samples were collected on two experimental voyages designed specifically to compare BW-associated biota before and after management. Microbial community structure and inferred function were assessed based on high throughput sequencing of 16S rRNA amplicons, and bacterial indicator taxa *E. coli* and enterococci were analyzed using targeted qPCR. As expected, both BWT alone and BWT+E dramatically changed bacterial communities, with the latter resulting in the largest overall decreases in bacterial diversity. Increases in Gammaproteobacteria, especially in the genus *Pseudomonas*, were particularly notable, with concomitant decreases in Alphaproteobacteria and Bacteroidia. Shifts in predicted bacterial function associated with BWT were similar for both voyages, despite significant differences in community structure, and may represent selection for *r*-strategists capable of active regrowth after BWT. qPCR estimates of indicator taxa were similar to those obtained through standard culture methods but may offer increased sensitivity for detecting changes associated with management. Our results indicate that BW management is effective at reducing bacterial communities but suggest that further research is needed to understand risks associated with taxa that may survive BWT.

## 1. Introduction

The translocation of living organisms in ballast water (BW) is a significant driver of global biotic exchange in aquatic environments and has been responsible for numerous biological invasions with devastating negative impacts on recipient systems [1-4]. International recognition of these risks has driven the development and implementation of various legal frameworks aimed at curtailing BW-borne invasions, most prominently the International Maritime Organization’s (IMO) 2004 International Convention for the Control and Management of Ship’s Ballast Water and Sediments [5], which entered into force in September 2017 [6]. Regulation D-2 of that Convention establishes numerical discharge limits for multiple size classes of BW-borne organisms, limits that have been similarly adopted in other regulations such as those established by the U. S. Coast Guard [7] and U. S. Environmental Protection Agency [8].

Achievement of discharge limits is anticipated to come primarily through technology-based shipboard treatment of BW (ballast water treatment, BWT) to reduce the concentration of living or viable organisms discharged into recipient systems [9]. Most international vessels were expected to have installed ballast water management systems (BWMS) by 2024 [10]; however, recent studies suggest that challenges remain to achieving full effectiveness of these systems in real world settings [11-13]. In light of these challenges, some vessels may still manage BW at least in part *via* BW exchange (BWE), the practice of flushing ballast tanks with open ocean water to decrease abundance and viability of coastal organisms [14]. BWE has been shown to be an effective strategy for mitigating introduction risk when BWMS are entirely or partially inoperable [13]. The combination of treatment and exchange (BWT+E) may be particularly protective when uptake concentrations are high and/or when BW is discharged into freshwater recipient systems [13, 15, 16]. Both modeling and empirical studies have shown that BWT+E would likely lower rates of invasion of non-native zooplankton in freshwater ports by reducing relative abundance of higher risk freshwater/euryhaline taxa and replacing them with marine taxa with lower risk of establishment [15, 16]. At least one study has shown that BWT+E may be more effective than BWT alone at reducing organismal abundance across all regulated size classes [17].

Understanding introduction risks posed by BW discharge and the effectiveness of management actions aimed at reducing those risks entails accurate assessment of biodiversity present in BW tanks and released into recipient systems. While much early research conducted to understand biodiversity transported with BW generally focused on macro-organisms [18-20] due in part to the inherent challenges of discriminating microbial diversity [21-23], the potential for microbial invasions has been appreciated for over two decades [24, 25]. Concerns have focused primarily on the spread of pathogens and other harmful species. BW harbors *Vibrio cholerae* [26, 27], *Enterococcus* spp. [28], as well as coral and fish pathogens [29, 30], among others, revealing risks associated with vectoring of pathogenic microbes through ballast [31]. This risk is specifically acknowledged in regulatory standards setting numerical discharge limits for pathogenic taxa; D-2 regulations set allowable limits of toxicogenic *V. cholerae, E. coli*, and intestinal enterococci to 1, 250, and 100 colony forming units (CFU) per 100 mL, respectively. Modern tools, such as High Throughput Sequencing (HTS), have revealed that free-living microorganisms exhibit distinct biogeographic patterns [32]. Heightened concerns that microbial invasions— beyond those associated with pathogenic species— may be commonplace and impactful [32-34] have stimulated increased interest in broader assessment of microbial communities in BW [35-37]. The degree to which introduced microbial diversity may have altered the structure and function of recipient ecological systems remains underappreciated, as does the potential for such introductions *via* BW and other vectors [38].

Reduction in the risk of such impacts is one of the major goals of BW management and BWT in particular. Nevertheless, it is widely recognized that disinfection of water may lead not only to overall reduction in bacterial abundance and diversity, but also to shifts in composition of bacterial communities with potentially unintended consequences. These effects have been observed in wastewater and drinking water treatment systems, where survival of particular taxa and subsequent regrowth have been shown to drive large changes in microbial assemblages [39], in some cases including selection for opportunistic pathogens [40] or taxa harboring antibiotic resistance genes (ARGs) [41]. These selective processes are known to operate during treatment of BW, as well. While BWMS have been shown to be effective at disinfecting BW, post-treatment regrowth of resistant taxa is also likely to result in observable overall shifts in bacterial community structure and function [38, 42-44]. Such alterations may have implications for risks associated with treated BW. For instance, studies have shown that reductions in overall abundance and diversity of BW may result in selection for taxa resistant to challenging environmental conditions, which may present heightened risk for invasion [45]. The presence in ballast tanks of bacteria known to harbor ARGs and exhibit resistance to disinfection processes further elevates concerns that BWT may not fully eliminate such risk. These results are consistent with theory predicting that selection during transport may be an important factor determining the risks posed by BW transport [46].

In the current study we use molecular methods to assess changes in BW-borne bacterial communities associated with BW management. We take advantage of a unique ship-board experiment allowing comparison of microbial communities pre- and post-management, including BWT alone as well as BWT+E. We utilize 16S rRNA-based amplicon sequencing to explore changes in overall bacterial community structure and employ analyses enabling assessment of shifts in functional diversity. In addition, we adopt standard molecular approaches to evaluate the effectiveness of BW management in reducing abundances of the indicator taxa *E. coli* and enterococci. Our aims are to: 1) demonstrate the potential value of molecular tools for assessing BW risk and the efficacy of management practices; 2) determine the degree to which BWT alone and BWT+E reduce overall bacterial abundance and diversity as well as the abundance of indicator taxa; and 3) to explore the possibility of post-management shifts in community structure and/or function that may provide important considerations for future risk assessment.

### 2.1 Sample collection

Representative samples were collected through a custom sampling device attached to the main ballast line. The device and sampling methods are described elsewhere [47] but briefly, whole water samples used here were supplied at a rate of ∼0.4 L per minute over an hour of sampling. Sampling occurred during the ballasting (uptake of ballast water, designated “UP”) and deballasting (discharge of ballast water, “DIS”) operations. Because the sample probe was placed upstream of the BWMS, sample water from the uptake was not filtered or treated by chlorination. The upstream sample therefore represents the community of organisms in the source water (Fig. 1A, top panel). For discharge sampling, samples were collected from water after filtration and chlorination, and after a period of sequestration (∼5 d) in the ballast tank (Fig. 1A, bottom panel). The DIS samples, however, were not dechlorinated, as this process occurs downstream of the sample port as water was discharged.

**Figure 1.**
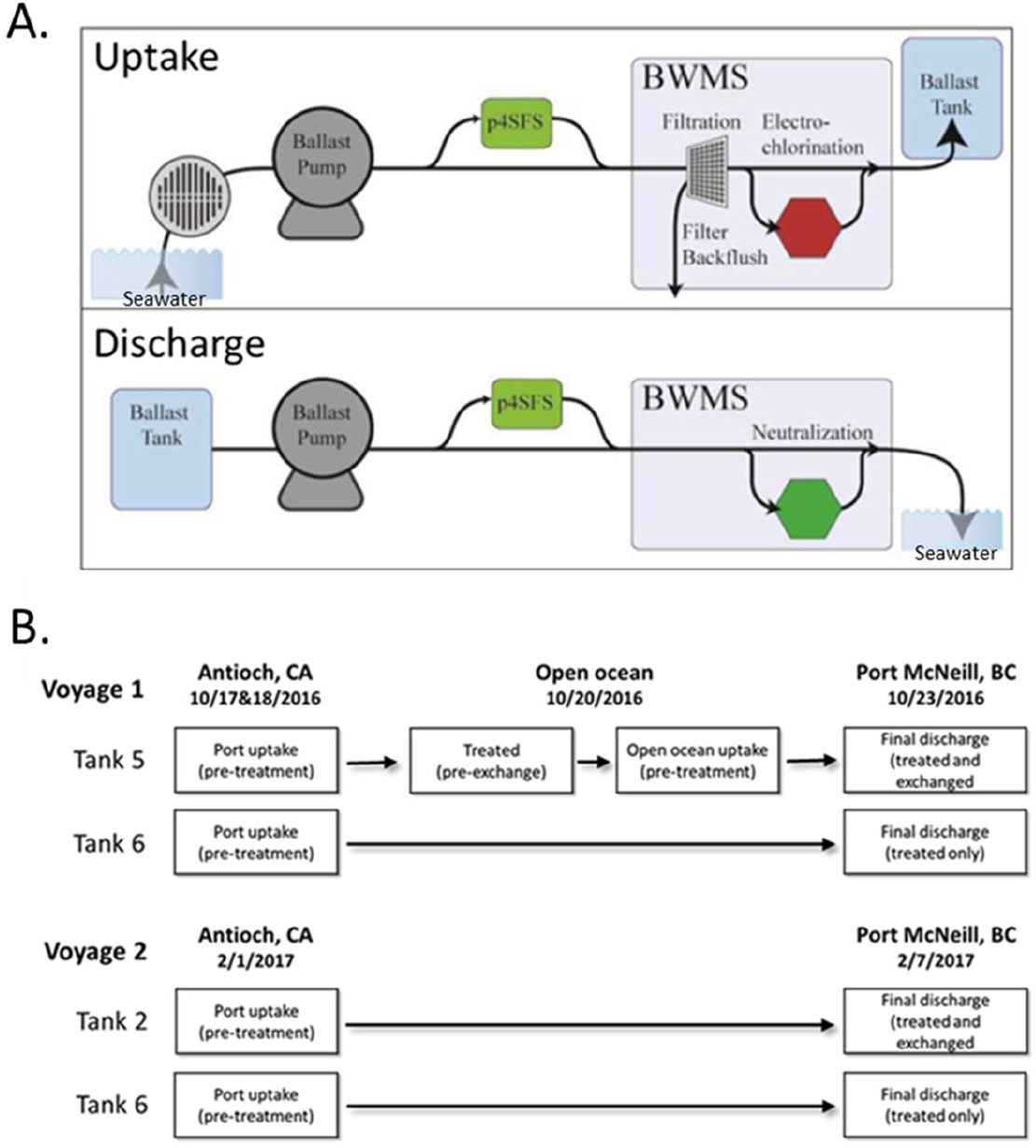
A. Schematic representation of ballast water management and sampling. p4SFS, NRL sampling skid; BWMS, ballast water management system. B. Sampling events on the two experimental voyages. Text in squares describes samples used in the analyses presented. Samples associated with open ocean exchange were not taken on Voyage 2.

A 15-mm diameter nominal (½” Nominal Pipe Size [NPS]) sample probe—inline to the p4SFS’s 50 mm diameter nominal (2” NPS) pipe—diverted a portion of the total sample flow through flexible tubing leading to a pre-cleaned, 20-L carboy. This sample (the “drip sample”) was collected at a low-flow rate into the carboy so a volume of 12-15 L of whole (unfiltered) water was collected during each sampling period (∼30 min). Throughout the sampling event, whole water was collected though the drip sampler at a rate of 0.4 to 0.7 L/min. Upon completion of sampling, a 500 mL aliquot of this whole water sample was collected for 16S rRNA sequencing.

### 2.2. Molecular methods

Samples were processed immediately upon receipt in the laboratory. Samples (150 ml of the 500 ml collected) were filtered through 0.45 µm mixed cellulose ester membrane filters (PALL Corporation, Westborough, MA) and then washed with 25 ml of phosphate-buffered saline. Membrane filters were aseptically transferred to sterile petri plates (PALL) and stored at -80°C.

Genomic DNA of microbes collected on membrane filters was extracted using the DNeasy PowerWater Kit (Qiagen, Germantown, MD) as instructed by the manufacturer. Prior to DNA extraction, petri plates with membrane filters containing concentrated ship ballast water samples were thawed to room temperature. Membrane filters were then rolled and inserted into PowerWater DNA Bead Tubes using sterile forceps. DNA yield was measured using the Qubit dsDNA HS Assay Kit (Invitrogen, Waltham, MA) and Qubit 2.0 Fluorometer (Invitrogen). Eluted DNA was stored at -80°C.

The strategy of Kozich et al. [48] was implemented to amplify and sequence variable region V4 of the 16S rRNA gene in ship ballast water samples using the 515F (5’-GTGCCAGCMGCCGCGGTAA-3’) and 806R (5’-GGACTACHVGGGTWTCTAAT -3’) primers. Amplicons were generated in triplicate PCR reactions using the FastStart High Fidelity PCR System (Millipore Sigma, St. Louis, MO). Reactions contained 5 µl of 10X Reaction Buffer, 1 µl of DMSO, 1 µl 10 mM dNTPs, 2 µl each barcoded forward and reverse primers, 0.5 µl Enzyme Blend and 1 ng sample DNA in a total of 50 µl. PCR conditions were 95°C for 2 min, followed by 25 cycles of 95°C for 30 sec, 55°C for 30 sec and 72°C for 1 min with a final extension of 10 min at 72°C. Products were viewed using 2% E-Gel 48 Agarose Gels (Invitrogen) with E-Gel Mother and Daughter E-Base (Invitrogen). Replicate reactions were pooled and then purified and normalized using the SequalPrep Normalization Plate Kit (Invitrogen) as instructed by the manufacturer, using 2 wells per pooled reaction to increase sample yield. Normalized samples were pooled by volume to create a library and the library was quantified using the KAPA Illumina Library Quantification Kit (Roche, Indianapolis, IN) and Agilent High Sensitivity DNA Kit (Agilent Technologies, Santa Clara, CA) as instructed by the manufacturer. The library was diluted to 3.5 pM, supplemented with 30% (v/v) PhiX Control v3 library (Illumina, San Diego, CA) and sequenced using the MiSeq Reagent Kit v2 (500 cycles, Illumina) and MiSeq instrument. Quality control reactions were amplified and sequenced along with samples to assess sequencing error and monitor for PCR contamination. Negative quality controls consisted of 10 mM Tris-HCl (pH 8.5) used for sample diluent. Positive quality controls were comprised of genomic DNA of 10 bacterial species mixed together at equivalent mass concentrations. The mock microbiome included *Streptococcus pneumoniae* (American Type Culture Collection (ATCC) #BAA334D), *Staphylococcus aureus* (ATCC #BAA-1718D), *Porphyromonas gingivalis* (ATCC #33277D), *Neisseria meningitidis* (ATCC #BAA-335D), *Listeria monocytogenes* (ATCC #BAA-679D), *Lactobacillus gasseri* (ATCC #33323D), *Deinococcus radiodurans* (ATCC #13939D), *Acinetobacter baumannii* (ATCC #17978D), *Bacillus cereus* (ATCC #10987D), and *Rhodobacter sphaeroides* (ATCC #17023D).

The 23S rRNA genes of enterococci and *E. coli* were quantified by qPCR as previously described [49, 50]. Briefly, each sample was analyzed in duplicate 25 µl PCR reactions that consisted of 1 µM each respective forward (enterococci: GAGAAATTCCAAAC-GAACTTG; *E. coli*: GGTAGAGCACTGTTTTGGCA) and reverse (enterococci: CAG-TGCTCTACCTCCATCATT; *E. coli*: TGTCTCCCGTGATAACTTTCTC) primers, 80 nM appropriate TAMRA probe (enterococci: TGGTTCTCTCCGAAATAGCTTTAGGGCTA; *E. coli*: TCATCCCGACTTACCAACCCG), 5 µg Bovine serum albumin (Millipore Sigma), TaqMan Environmental Master Mix (ThermoFisher Scientific, Lenexa, KS), PCR-grade water and 5 µl of sample. The standards consisted of a synthetic DNA fragment that included both the enterococci and *E. coli* amplicons and were run along with the samples at 6 dilutions ranging from 25845.9 – 5.7 copies per reaction. A StepOnePlus Real-Time PCR System (ThermoFisher Scientific) was used to perform PCR where conditions were 95°C for 10 min followed by 40 cycles of 95°C for 15 sec and 60°C (56°C for *E. coli*) for 1 min. Cycle threshold values were determined after using the AUTO feature in the software to determine baseline cycles and manually adjusting the threshold to 0.03 ΔRN. Quantities of 23S rRNA gene copies per 100 ml of samples were determined by interpolating the cycle threshold values of the standard curve and adjusting for the original volume of sample filtered. Mean genome equivalents (MGE, a molecular analogue for cell counts) were determined by dividing 23S rRNA gene copies by 7 for *E. coli* and 4 for enterococci [49, 50].

### 2.3. Bioinformatics

Demultiplexed paired-end FASTQ files were imported to QIIME2 (version 2022.11; [51]). Using the DADA2 QIIME2 plugin (version 1.14) [52], reads from demultiplexed FASTQ files were trimmed on their 3’ end to remove the primers. Quality filtering and denoising were performed using the default DADA2 parameters, and reads were merged using a default minimum overlap of 12 bp. Chimera removal was performed using the consensus method, which discards sequences found to be chimeric in the majority of samples. To assign taxonomy to sequences, a Naïve Bayes classifier (q2-feature-classifier; [53]) was trained on sequences with the same primers acquired from the SILVA database (version 138.1; [54-56]) using RESCRIPt (Robeson et al., 2021). Results from QIIME2 were imported into RStudio (version 4.4.0) using the qiime2R package (version 0.99.6), and non-target sequences, including those assigned to Archaea, Chloroplasts, and Mitochondria, were removed using the phyloseq package (version 1.48.0; [57]). Prior to conducting further analyses in phyloseq, the sequencing depth per sample was visualized with the rarecurve function in vegan (version 2.5.6; [58]) and samples with less than 1000 reads were removed from the dataset (Fig. S1). As rarefaction curves plateaued for all samples, suggesting complete amplicon sequence variant coverage, no data rarefaction was performed on downstream analysis.

### 2.4. Data analysis

Samples were grouped by voyage and sample type (i.e., port uptake, BWT, ocean uptake, BWT+E) and the estimate-richness function in the phyloseq package was used to calculate the Shannon diversity and richness of each group of samples. Statistical significance was evaluated using pairwise Wilcoxon Rank Sum tests (stats package, version 4.4.0). A principal coordinate analysis (PCoA) based on the Bray-Curtis dissimilarity matrix was conducted on relative abundance transformed data using the ordinate function in the phyloseq package, and PERMANOVAs were performed using the adonis2 function in the vegan package in order to determine the statistical significance of differences between groups. Bar charts examining taxonomic diversity were constructed using the fantaxtic package (version 0.2.0).

Predictive functional annotation was conducted using the software Phylogenetic Investigation of Communities by Reconstruction of Unobserved States (PICRUSt2), which uses hidden state prediction methods to infer functional potential from 16S rRNA amplicon sequence data [59]. Prior to functional annotation, archaea, chloroplasts, mitochondria, control samples, and samples containing less than 1,000 reads were removed from the dataset. Pathway-level inference and functional annotation was conducted using the MetaCyc database [60], and metabolic pathways generated by PICRUSt2 were imported into R for further analysis. To identify functions that increased in abundance after BWT, a differential abundance analysis was conducted using the DESeq2 package (version 1.44.0; [60, 61]). Due to limited BWT+E samples and the constraints of DESeq2, the functional potential of BWT+E communities could not be compared to that of BWT or ocean uptake samples using DESeq2; instead, differential abundance analyses have been limited to comparisons of functional potential between port uptake and BWT samples from each voyage. In order to compare shifts in functional potential to shifts in bacterial community composition, metabolic pathways were log transformed and visualized using a PCoA plot based on the Bray-Curtis dissimilarity matrix.

Sequence data is available in the NCBI Sequence Read Archive (SRA) under the accession number PRJNA1282192; all code used to generate figures and conduct analyses is available on Github (https://github.com/sbrown2648/Ballast-water-bacteria).

## 3. Results

### 3.1. Bacterial diversity

Sequencing of the V4 region of the 16S rRNA gene yielded a total of 154,760 reads, which corresponded to 459 ASVs identified as bacteria. Of these ASVs, 17 were assigned species-level identification, and 265 were assigned genus-level identification. The number of reads varied between sample type (i.e., port uptake, BWT, ocean uptake, and BWT+E) and voyage (Fig. S2). BWT+E samples from voyage 2 exhibited read abundances below our threshold for inclusion and were left out of downstream analyses.

The richness and evenness of bacterial communities differed with sample type and voyage. Mean values of bacterial richness, or the number of observed ASVs, decreased after treatment in ballast water from both voyages, with the average number of unique ASVs decreasing from 41 to 28 post-treatment in ballast water from voyage 1, and from 71 to 27 in post-treatment ballast water from voyage 2 (Fig. 2a). However, this decrease was only significant in samples collected from voyage 2 (Pairwise Wilcoxon Rank Sum Test, *p* = 0.002). In ballast water that underwent both treatment and exchange, the richness of bacterial communities was lower than in ballast water that underwent treatment alone (19 ASVs on average), though this difference was not statistically significant (Pairwise Wilcoxon Rank Sum Test, *p* = 0.082). In contrast, Shannon diversity values were similar between BWT and BWT+E samples (2.03 and 2.04, respectively; Pairwise Wilcoxon Rank Sum Test, *p* = 0.840) (Fig. 2b).

**Figure 2.**
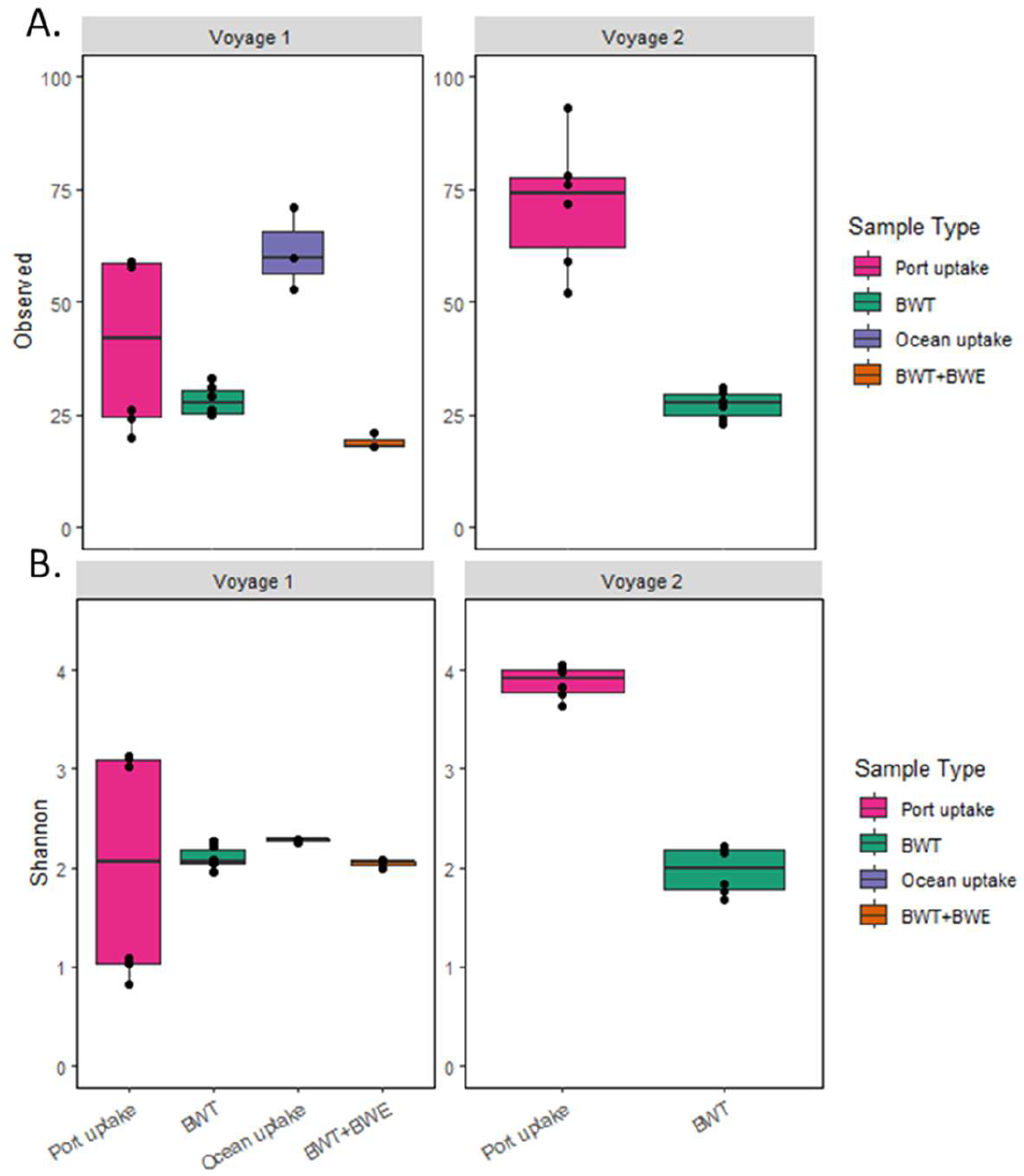
Comparison of diversity between sample types. A. Observed richness and B. Shannon diversity. Sample types across multiple tanks were combined for analyses; green BWT boxes for voyage 1 include both BWT only samples from tank 6 and pre-exchange BWT samples from tank 5.

### 3.2. Changes in microbial community structure

Significant changes in bacterial community composition occurred in ballast water after BWT and BWT+E (Table S1). After BWT and BWT+E, bacterial communities were dominated by members of the genus *Pseudomonas* that were initially present in relatively low abundances in port and ocean uptake samples (Fig. 3). During voyage 1, individual ballast water tanks exhibited similar changes in bacterial community composition (Fig. S3): bacteria in the genus *Pseudomonas*, which initially comprised an average of 10% of the relative abundance of port uptake samples increased to 29% of the total relative abundance after BWT (Fig. 3). Notably, ASVs identified as the fish pathogen *Pseudomonas anguilliseptica* accounted for 19% of the total relative abundance in BWT samples from voyage 1, a substantial increase from their initial abundance of 6% in port uptake samples (Fig. 3). Bacterial communities in ocean uptake samples were distinct from other sample types, and primarily dominated by members of the Gammaproteobacteria, including bacteria in the Vibrionaceae family and the genera *Shewanella, Pseudomonas, Pseudoalteromonas*, and *Oleispira* (Fig. 3, Fig. 4a, Table S1). In samples discharged after BWT+E, the relative abundance of *P. anguilliseptica* had decreased to 0.9% of the total relative abundance, though bacteria in the *Pseudomonas* and *Pseudoalteromonas* genera remained dominant in these samples (Fig. 3).

**Figure 3.**
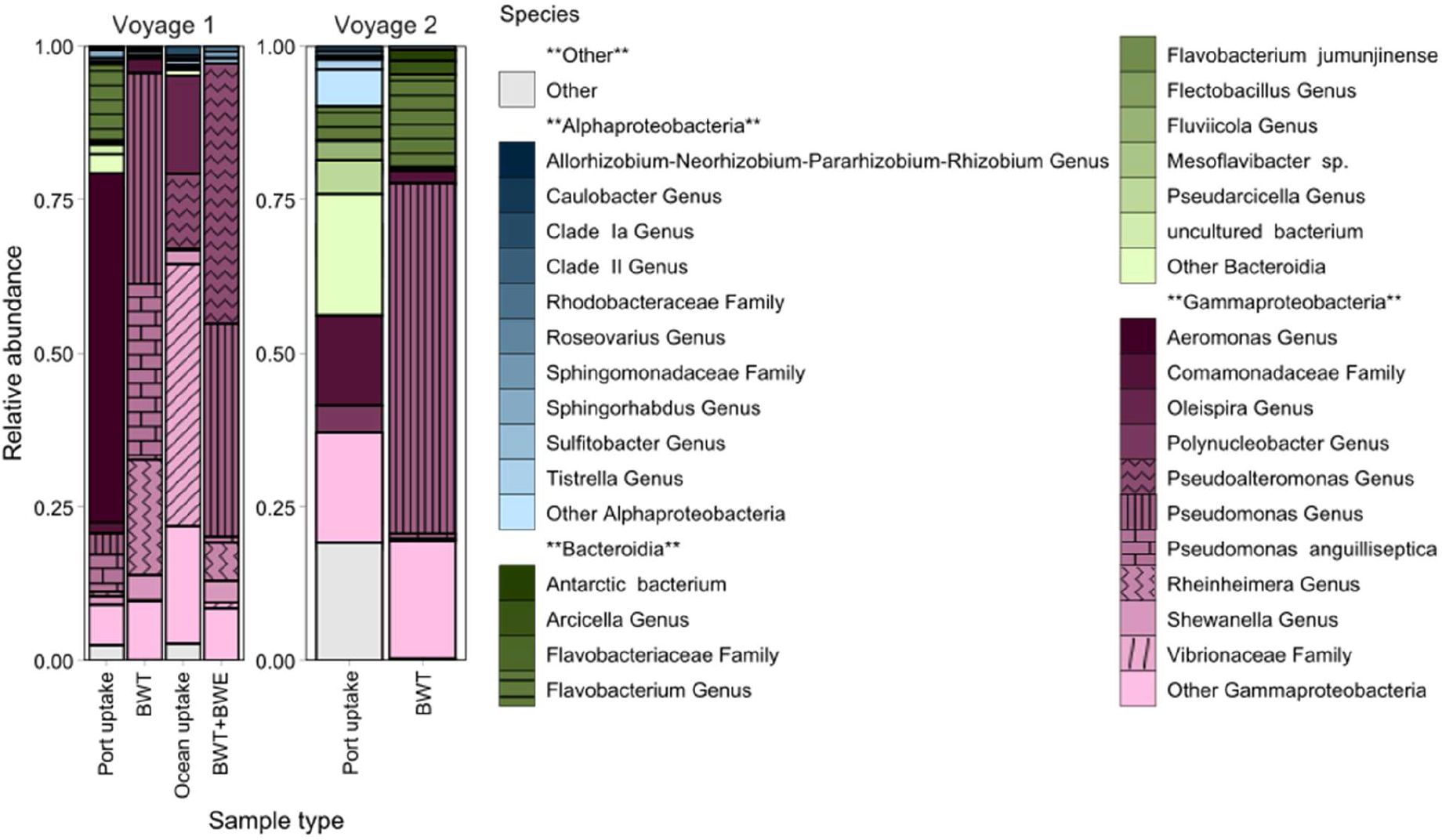
Bar plot showing the relative abundance of the top 10 most abundant species-level identifications observed within the top three classes across sample types. BWT+E samples from voyage 2 exhibit read abundance too low to analyze. Blue shading indicates taxa within the class Alphaproteobacteria, green indicates Bacteroidia, red indicates Gammaproteobacteria, and grey indicates other classes of bacteria.

**Figure 4.**
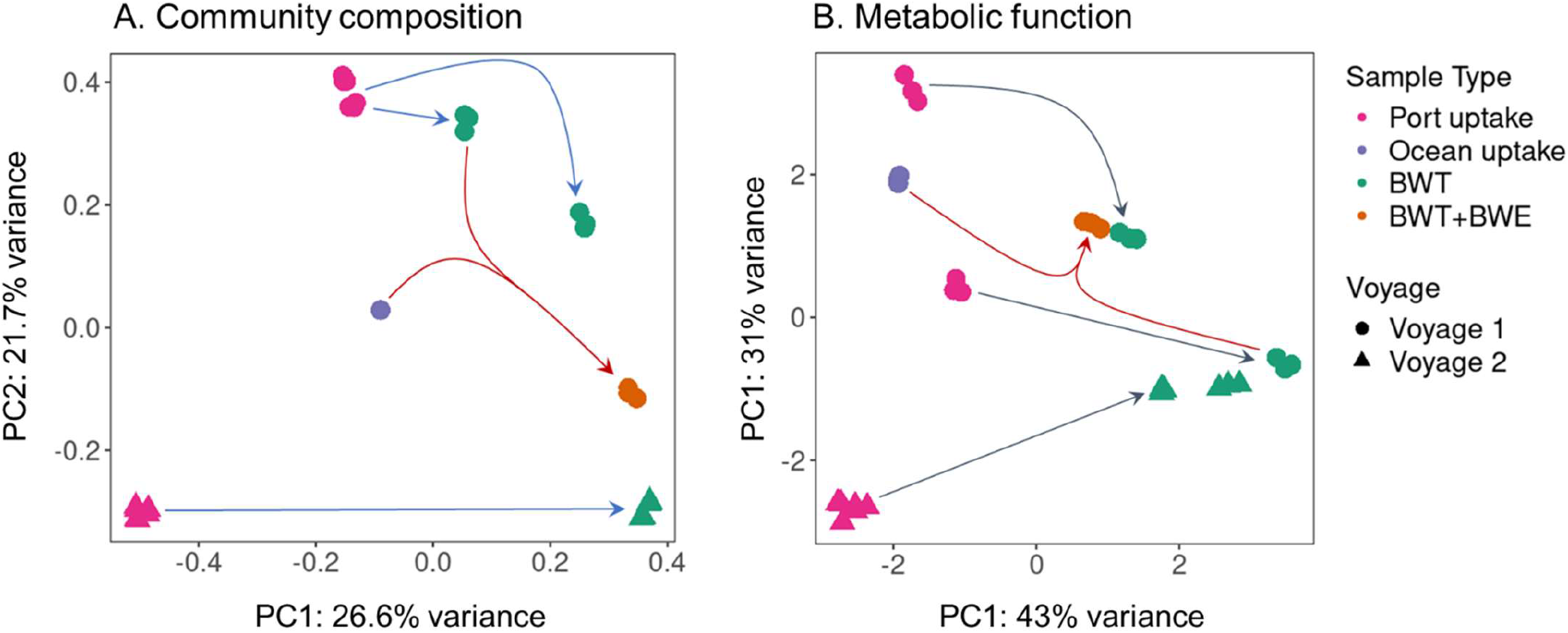
Ordination of samples based on 16S community composition (A) and metabolic function determined by PICRUST analysis (B). Arrows connect samples before and after management actions; blue lines indicate BWT, red lines indicate BWT+E and represent addition of ocean uptake to treated BW.

Community composition in BWT samples from voyage 2, while distinct from that of voyage 1 (Fig. 4a) shifted in a similar manner after ballast water treatment, with *Pseudomonas* becoming the most abundant genus (57% of the total relative abundance) (Fig. 3). However, *Pseudomonas anguilliseptica* was present in much lower relative abundances (0.7%) in BWT samples from voyage 2 than in those from voyage 1 (Fig. 3). Notably, BWT samples from voyage 2 had higher proportions of bacteria in the Bacteroidia class, including the genera *Flavobacterium* and *Arcicella*.

### 3.3. Changes in microbial community metabolic function

A differential abundance analysis identified 92 inferred metabolic pathways that significantly increased in abundance in ballast water samples after treatment, including pathways corresponding to central metabolic functions (TCA cycle, glyoxylate cycle), biosynthesis of lipids, amino acids, and secondary metabolites, and aromatic compound degradation (Fig. S4). The functions with the greatest increase in abundance (>1 log2Fold-Change) in ballast water after treatment included pathways related to the degradation of amines and polyamines, amino acids, aromatic compounds, and carbohydrates (Fig. 5). Additionally, pathways associated with fermentation, methylphosphonate degradation, the TCA cycle, and the biosynthesis of cofactors, prosthetic groups, electron carriers, and vitamins had some of the greatest positive log2FoldChanges (Fig. 5). Notably, shifts in metabolic function after treatment exhibited similar patterns in tanks from both voyages, and predicted metabolic functions in BWT and BWT+E samples remained distinct from port and ocean uptake communities (Fig. 4b).

**Figure 5.**
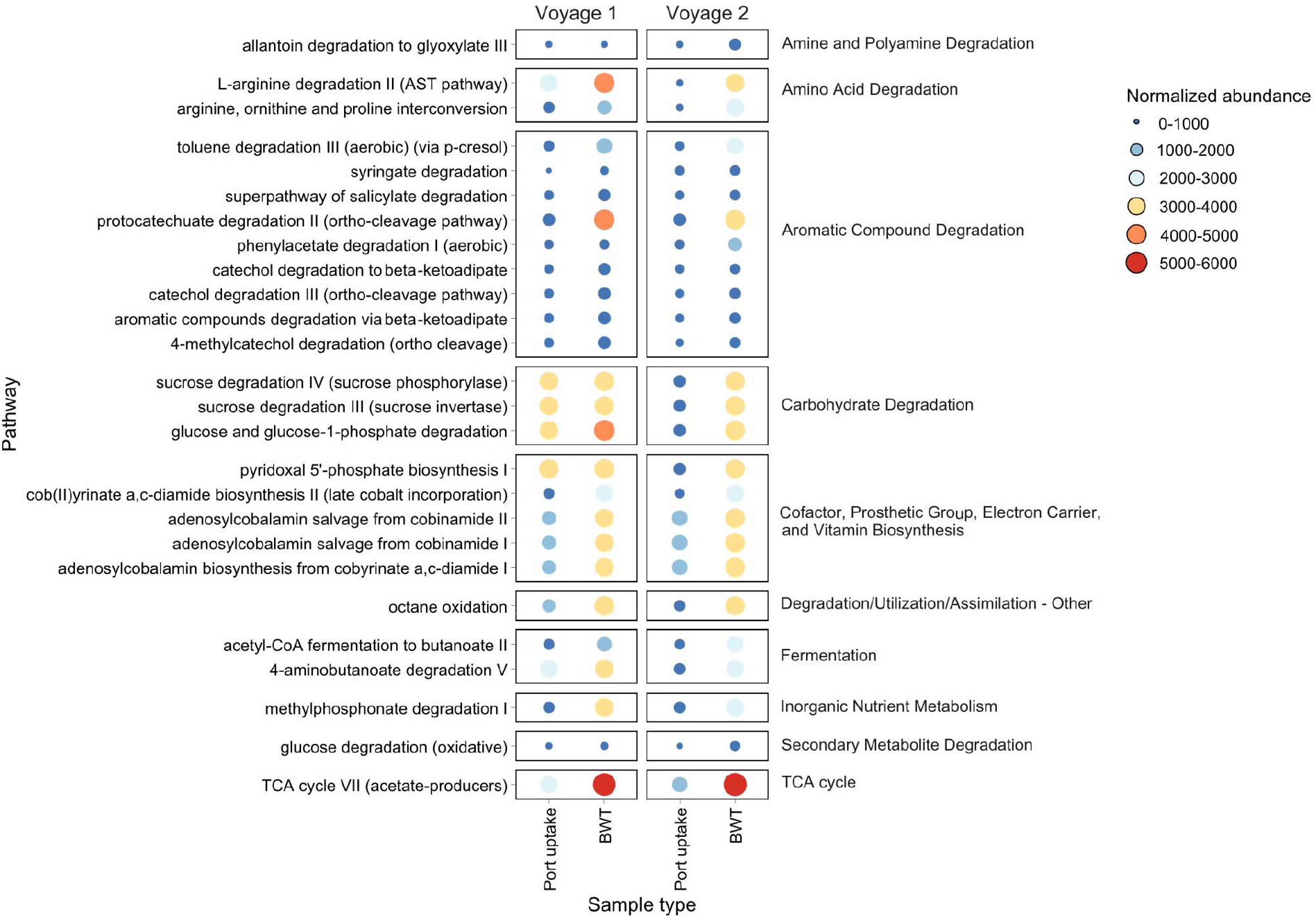
The normalized abundance of inferred metabolic pathways with a log2FoldChange greater than or equal to 1 in BWT compared to port uptake.

### 3.4. Comparison of qPCR and MPN methods for indicator taxa

BW management (either BWT or BWT+E) consistently resulted in decreases in indicator cell counts as estimated by qPCR; in fact, for enterococci all tests showed reductions to undetected MGE per 100 mL (Table 1). Overall *E. coli* estimated counts decreased from 36.4 ± 15.1 to 7.7 ± 10.9 (mean + standard deviation; Student’s t-test P = 0.0005) and enterococci decreased from 29.5 ± 21.3 to below the limit of detection (P = 0.0012). In only one test (Voyage 1, T5) did the estimate for *E. coli* MGE increase; despite registering below the limit of detection after initial BWT and below detection limits in open ocean water, the discharged sample after BWT+E detected the presence of E. coli gene fragments but was below the limits of quantification (5.7 MGE/100 mL). The MPN approach (results reported previously in [62] and reproduced here) also consistently showed decreases in estimated counts for both indicators, although it appeared to be generally less sensitive to changes associated with management, in part because many tests were limited to a threshold of 10 cells/100 mL. All samples tested, including uptake samples prior to BW management, were estimated to be in compliance with the D-2 discharge standard for both indicators, regardless of method.

**Table 1.**
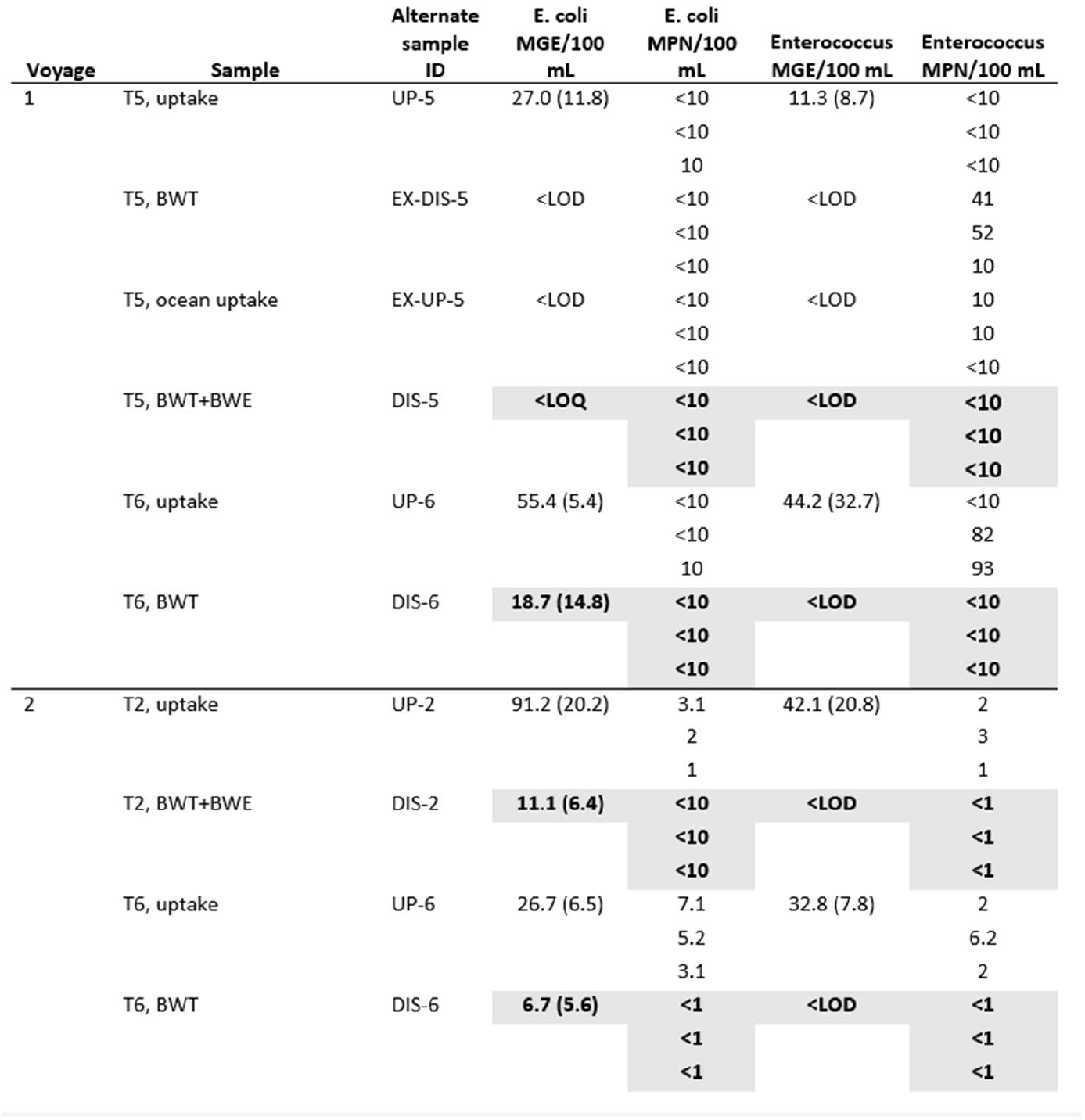
Comparison of most probable number (MPN) and qPCR estimates for indicators *E. coli* and enterococcus. MGE = mean genome equivalents. ‘Sample’ column includes the tank ID and the type of sample. Shading indicates port discharge samples that comply with US and IMO standards (<250 colony forming units/100 mL for *E. coli*, <100 colony forming units/100 mL for enterococcus). Results of three MPN trials are listed for each sample. For qPCR results, the mean of three samples is shown with standard deviations in parentheses; the Limit of Detection (LOD) for *E. coli* is 1.9 MGE/100 mL and enterococci is 3.3 MGE/100 mL. The limit of quantification (LOQ) is 5.7 MGE/100 mL for *E. coli* and 10 MGE for enterococci.

## 4. Discussion

Few studies have directly assessed changes to bacterial communities associated with BW management; even fewer have been able to evaluate differences in bacterial diversity before and after management on real voyages. However, existing research does consistently suggest that 1) both BWT [42-44, 63] and BWE [35] substantially reduce and alter, but do not eliminate, bacterial communities; 2) BWT typically results in predictable shifts in community structure away from Alphaproteobacteria and towards Gammaproteobacteria [44, 63]; and 3) bacterial communities exhibit the capacity to regrow after BWT, with *r*-strategists becoming more dominant [42-44]. Our results, derived from unique experimental voyages allowing direct comparisons of pre- and post-management diversity in real operational contexts, are consistent with these findings and provide important additional insights into the effects of BW management on bacterial communities.

Overall bacterial richness and diversity declined with management during both voyages, in agreement with previous studies [42, 63], though declines in diversity were not always statistically significant. The most dramatic declines were observed for voyage 2, where significant reductions in both richness and diversity occurred with BWT (Figure 2). While declines were also observed for voyage 1, they were less dramatic. This difference in response to management may be, in part, a function of differences in the bacterial diversity present in uptake samples between the two voyages. Voyage 1 uptake exhibited much lower diversity than voyage 2 uptake; while post-BWT discharge samples for both voyages had similar overall diversity, declines observed for voyage 1 were less pronounced due to lower pre-BWT diversity. Interestingly, BWE results in an apparent influx of bacterial diversity from the open ocean, such that the overall modest decline in diversity of BWT+E relative to BWT alone in voyage 1 actually reflects rather sizable reductions in diversity subsequent to this influx (Fig 2a and b). Unfortunately, we can only assess effects of BWT+E for that voyage, since sampling associated with mid-ocean exchange was not conducted on voyage 2 and the BWT+E samples taken on final discharge yielded too few sequences to be considered for downstream analyses. However, this may be an additional indication that BWT+E is effective in reducing bacterial concentrations and diversity.

Declines in bacterial community richness and diversity after BWT and BWT+E were accompanied by substantial shifts in community structure leading to bacterial communities distinct from those of natural seawater (here, port uptake and ocean uptake communities), as documented by previous studies (Fig. 4A) [44, 63, 64]. General shifts away from Alphaproteobacteria and Bacteriodia and towards Gammaproteobacteria were observed for both voyages and for both BWT alone and BWT+E (Figure 3). Such shifts at higher taxonomic levels have been observed in other studies, suggesting that selective recolonization of *Gammaproteobacteria* occurs in response to multiple BWT modalities [44, 63]. Perhaps most notable were dramatic increases in genera containing potentially pathogenic bacteria – in BWT and BWT+E, bacteria in the *Pseudomonas* genus became highly abundant, including those identified as *Pseudomonas anguilliseptica*, a fish pathogen, which accounted for 19% of the total relative abundance in BWT from voyage 1 (Fig. 3). Notably, BWT+E effectively reduced the abundance of *P. anguilliseptica* to 0.9%. However, members of the Pseudomonas genus, a group containing several notable pathogens, including *P. aeruginosa* and *P. maltophilia*, continued to comprise a substantial portion of the relative abundance in BWT+E samples (Fig. 3). Although 16S amplicon sequencing has several notable limitations when used for the detection of potentially pathogenic bacteria (e.g., a limited ability to resolve taxa to the species level or determine the pathogenicity risk of closely related strains [65]) 16S detections of potentially pathogenic bacteria in ballast water have been found to be similarly detected by qPCR [66]. 16S rRNA amplicon sequencing may also be an effective tool for use in pre-screening bacterial communities prior to qPCR, in order to select potentially pathogenic genera to focus qPCR species-level detections on [67]. Additionally, qPCR has been shown to possess a greater capability for detecting pathogens present in low relative abundance that go undetected by 16S rRNA amplicon sequencing, suggesting that sequencing results may be underestimating the number of pathogens present in ballast water [66, 67]. A combination of 16S rRNA amplicon sequencing and qPCR, as previously suggested by Lv et al. and Cui et al. [66, 68], may thus be the most effective method for assessing the presence of potential pathogens in ballast water.

Despite re-growth of bacterial communities post-BWT or BWT+E, estimates of bacterial community richness and evenness remained lower than pre-treatment estimates, demonstrating the known efficacy of BWT and BWT+E to reduce overall bacterial community diversity [43, 63]. While bacterial communities in ballast water experience substantial reductions in diversity after BWT, bacterial diversity tends to increase over the course of a voyage, with the duration of storage time post-BWT acting as a significant factor in determining the abundance and diversity of bacteria, including pathogenic bacterial species [43, 64]. However, our results indicated that storage time had little effect on bacterial diversity, as treated water held for 6 days (samples from tank 6 of voyage 1) and treated water held for only 2 days (samples from tank 5) exhibited nearly identical diversity measures (left-hand panels in Fig 2a and b, green boxes). This lack of notable differences in bacterial diversity despite increases in the length of storage time may be a result of the length of time tested. Alternatively, such shifts in diversity over time may be dependent upon the biogeography of the source water or other environmental factors [64]. These results highlight the complexity in accurately predicting bacterial community responses post-BWT or BWT+E, which are essential to understand in order to accurately determine the invasion risk these communities pose to recipient communities.

In addition to substantial changes in community structure, BWT and BWT+E resulted in changes in predicted bacterial community metabolic function, leading to post-BWT bacterial communities possessing a functional potential distinct from that of the original communities (port or ocean uptake) (Fig. 4b). A differential abundance analysis revealed that significant increases in inferred pathways associated with carbon metabolism, including the TCA cycle, amino acid degradation, and carbohydrate degradation, were expressed to a greater extent after BWT than in port uptake samples during both voyages, corresponding with increases in Gammaproteobacteria in BWT samples (Fig. 5, S4). These functions, in addition to being essential for bacterial survival, may indicate bacterial responses to increases in labile organic matter availability, which can occur as a result of cells being lysed or damaged during BWT [42, 43]. Due to limitations in sample size, we were unable to conduct a differential abundance analysis comparing functional changes in BWT+E to those in BWT or ocean uptake samples; however, a PCoA plot indicated that similar shifts in inferred functional capabilities in BWT+E occurred as in BWT, despite differences in bacterial community composition (Fig. 4). As the post-BWT and BWT+E bacterial communities exhibited unique functional capabilities, their introduction into non-native environments may risk altering bacterially-mediated ecosystem functions [38].

Our examination of indicator taxa represents a relatively weak test of the efficacy of BWT or BWT+E, as all uptake samples exhibited indicator counts below the D-2 standard. This is consistent with previous observations that much unmanaged BW achieves regulatory discharge limits even in the absence of treatment [69]. BWT appears to have been more effective in eliminating *Enterococcus* than *E. coli*, with all post-BWT counts for *Enterococcus* at levels below detection limits (for qPCR) or near zero (for MPN). Our results reveal a general consistency, both in terms of overall abundances and patterns of reduction associated with BW management, between a standard, commonly employed method of assessing indicator taxa in BW (MPN) and qPCR methods developed for use in other water quality contexts (primarily recreational freshwaters) [49, 50]. This suggests the potential value of such methods for BW monitoring and assessment.

Other studies have employed qPCR approaches to measure the presence of pathogenic taxa in BW; in one recent study, *E. coli* was observed in all BW samples examined, with gene copy numbers ranging from 10^3^ to 10^5^ per 100 mL [66]. Raw copy numbers observed in the current study were considerably lower, ranging from 187 to 638 per 100 mL in uptake samples. However, the qPCR methodologies utilized here were different and allowed for estimation of MGE (analogous to cell counts) and direct comparison to the standard MPN approach. Generally, the qPCR results reported here appear to exhibit greater sensitivity than MPN to changes in indicator cell counts associated with BW management, although some uncertainties remain and require further investigation. For instance, it is not clear whether the increase in *E. coli* MGE after BWT+E in Voyage 1 tank 5 reflects error in the method or, possibly, regrowth from levels below the qPCR detection limit during the sequestration period prior to discharge (Table 1).

## 5. Conclusions

Our examination of changes in bacterial community structure and function in response to BWT and BWT+E during experimental voyages provides additional insight into the effects of BW management on bacterial communities. Both BWT and BWT+E effectively reduced the concentrations of indicator bacteria in ballast water to levels below the D-2 standard, with qPCR detections of indicator bacteria providing greater sensitivity than the standard MPN approach. However, while both BWT and BWT+E effectively reduced bacterial diversity and the presence of indicator bacteria, 16S amplicon sequencing results detected the presence of potentially pathogenic bacteria in ballast water post-BWT and BWT+E, suggesting that treated or treated and exchanged ballast water may still pose a risk for pathogen introduction into recipient ecosystems. Further, BWT and BWT+E altered the inferred functional potential of ballast water bacterial communities such that post-BWT and BWT+E communities exhibited functional capabilities distinct from those of natural waters. Overall, the results of this study highlight the benefit in using a variety of molecular methods to examine bacterial communities present in ballast water, as well as the importance of effective BW management practices.

## Supplementary Materials

see associated file “Supplementary Tables & Figures.”

## Author Contributions

All authors contributed to conceptualization and planning of the work described. VM, MF and LD designed and led sampling efforts. SB and SPK generated and curated sequencing data; SB, SPK, OL, and AZ conducted formal analyses of sequencing data. NEB generated and analyzed qPCR data. SB and JAD wrote the initial draft. All authors reviewed and edited the final draft. All authors have read and agreed to the published version of the manuscript.

## Acknowledgments

This research was supported in part by an appointment to the U.S. Environmental Protection Agency (EPA) Research Participation Program administered by the Oak Ridge Institute for Science and Education (ORISE) through an interagency agreement between the U.S. Department of Energy (DOE) and the U.S. Environmental Protection Agency. ORISE is managed by ORAU under DOE contract number DE-SC0014664. The views expressed in this article are those of the authors and do not necessarily reflect the views or policies of the U.S. EPA, DOE, or ORAU/ORISE. We would like to thank Varun Rao for lab support and Mara Cuebas Irizarry and Eric Villegas for providing feedback on an earlier draft of the manuscript.

