## Supplemental figures and tables for "Molecular methods reveal responses of bacterial communities, including indicator species, to ballast water management"

Table S1. Results of a Pairwise PERMANOVA examining differences in bacterial community composition between sample types.

| pairs | Df | SumsOfSqs | F.Model | R2 | p.value | p.adjusted sig |
|---|---|---|---|---|---|---|
| Port uptake vs BWT | 1.00 | 2.44 | 8.88 | 0.29 | 0.00 | 0.0048 |
| Port uptake vs Ocean uptake | 1.00 | 1.70 | 6.52 | 0.33 | 0.00 | 0.0048 |
| Port uptake vs BWT+BWE | 1.00 | 1.63 | 6.22 | 0.32 | 0.00 | 0.0048 |
| BWT vs Ocean uptake | 1.00 | 1.82 | 8.80 | 0.40 | 0.00 | 0.0048 |
| BWT vs BWT+BWE | 1.00 | 0.97 | 4.65 | 0.26 | 0.00 | 0.0048 |
| Ocean uptake vs BWT+BWE | 1.00 | 1.26 | 107.32 | 0.96 | 0.10 | 0.1000 |

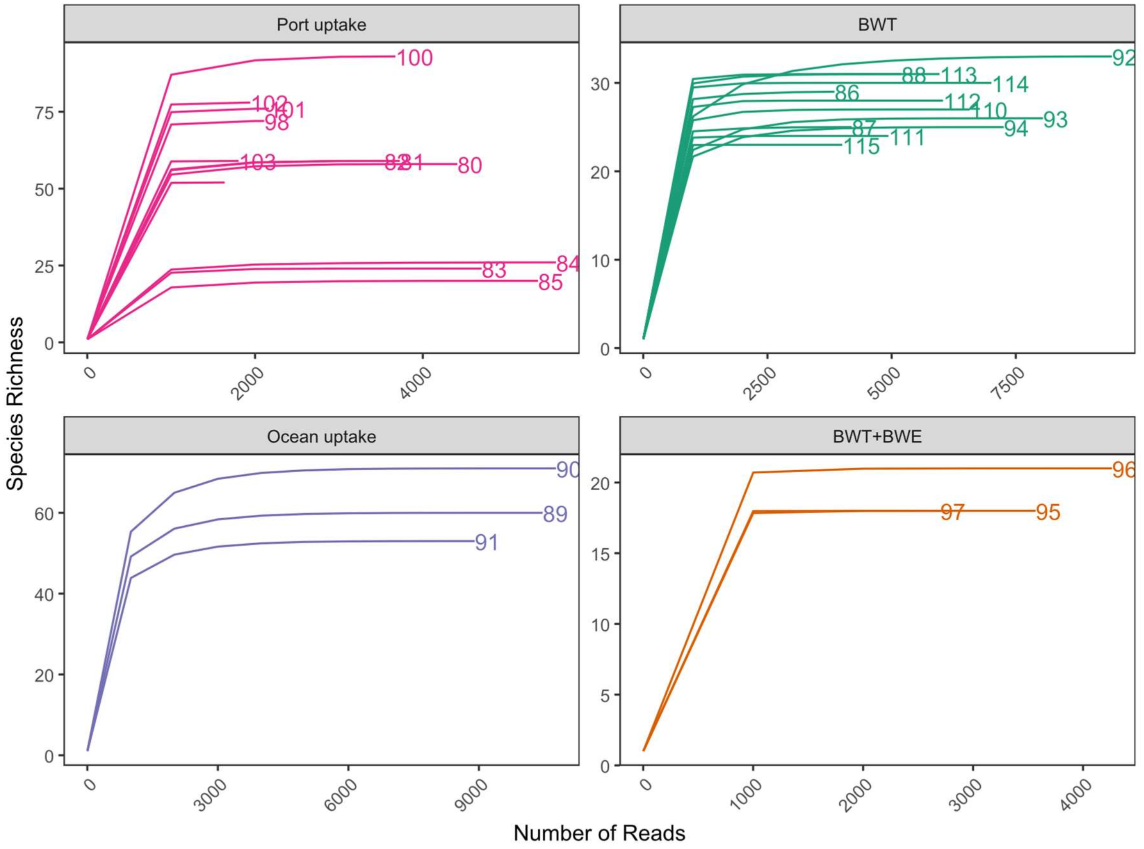


Figure S1. Rarefaction curves for each sample, separated by sample type. The sample numbers are shown next to each line. Samples with low numbers of reads (<1000) were removed prior to analysis.

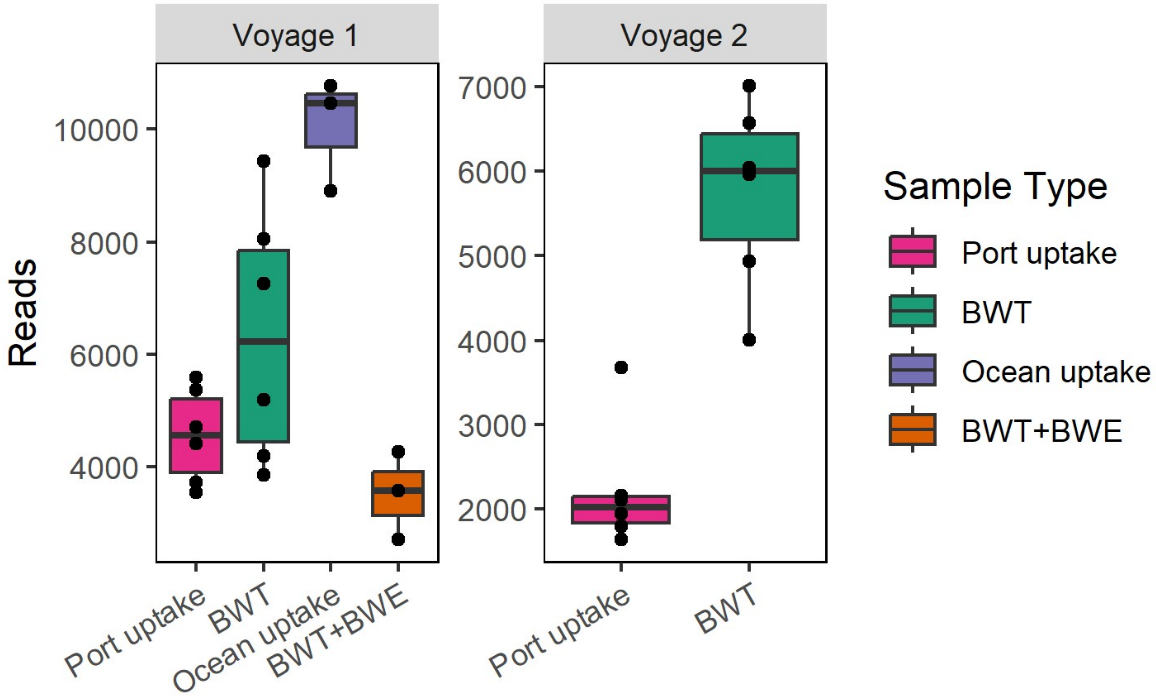


Figure S2. The number of reads per sample for each sample type and voyage.

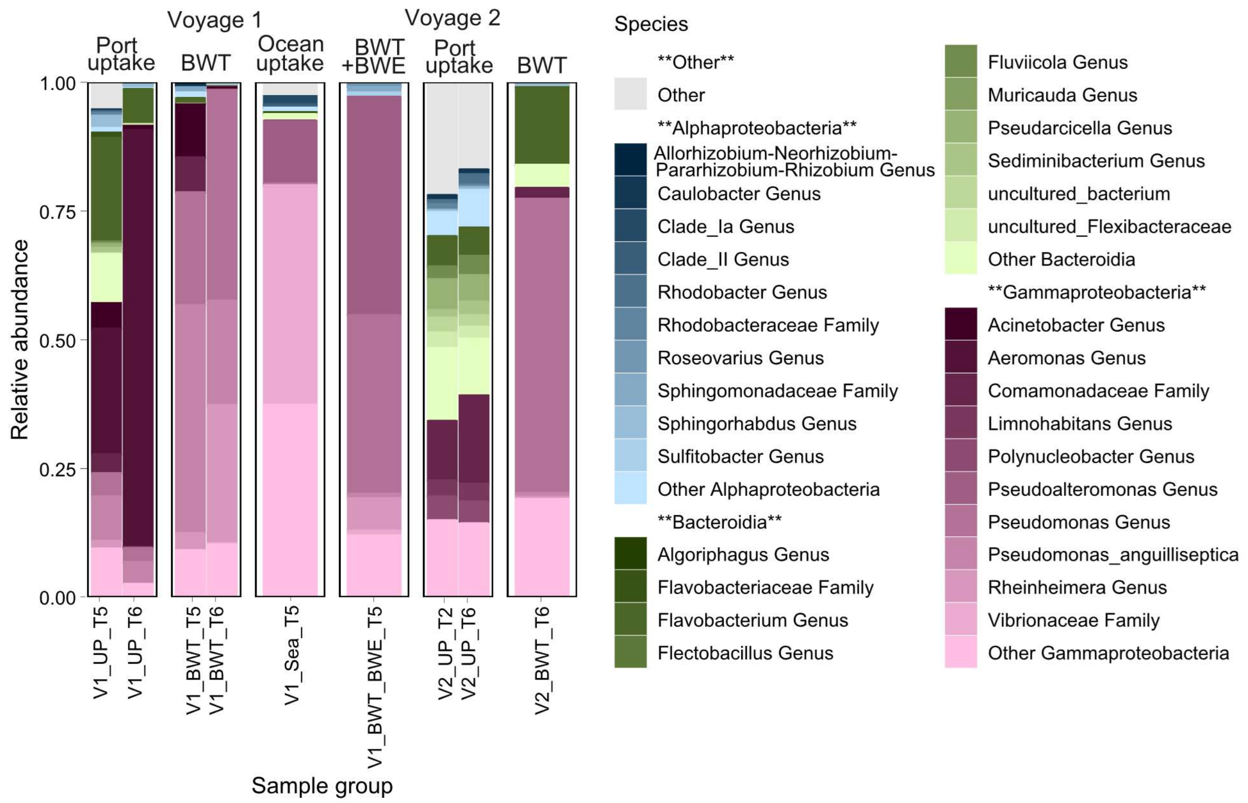


Figure S3. Bar plot showing the relative abundance of the top 10 most abundant species-level identifications observed within the top three classes across samples. Blue shading indicates taxa within the class Alphaproteobacteria, green indicates Bacteroidia, red indicates Gammaproteobacteria, and grey indicates other classes of bacteria.

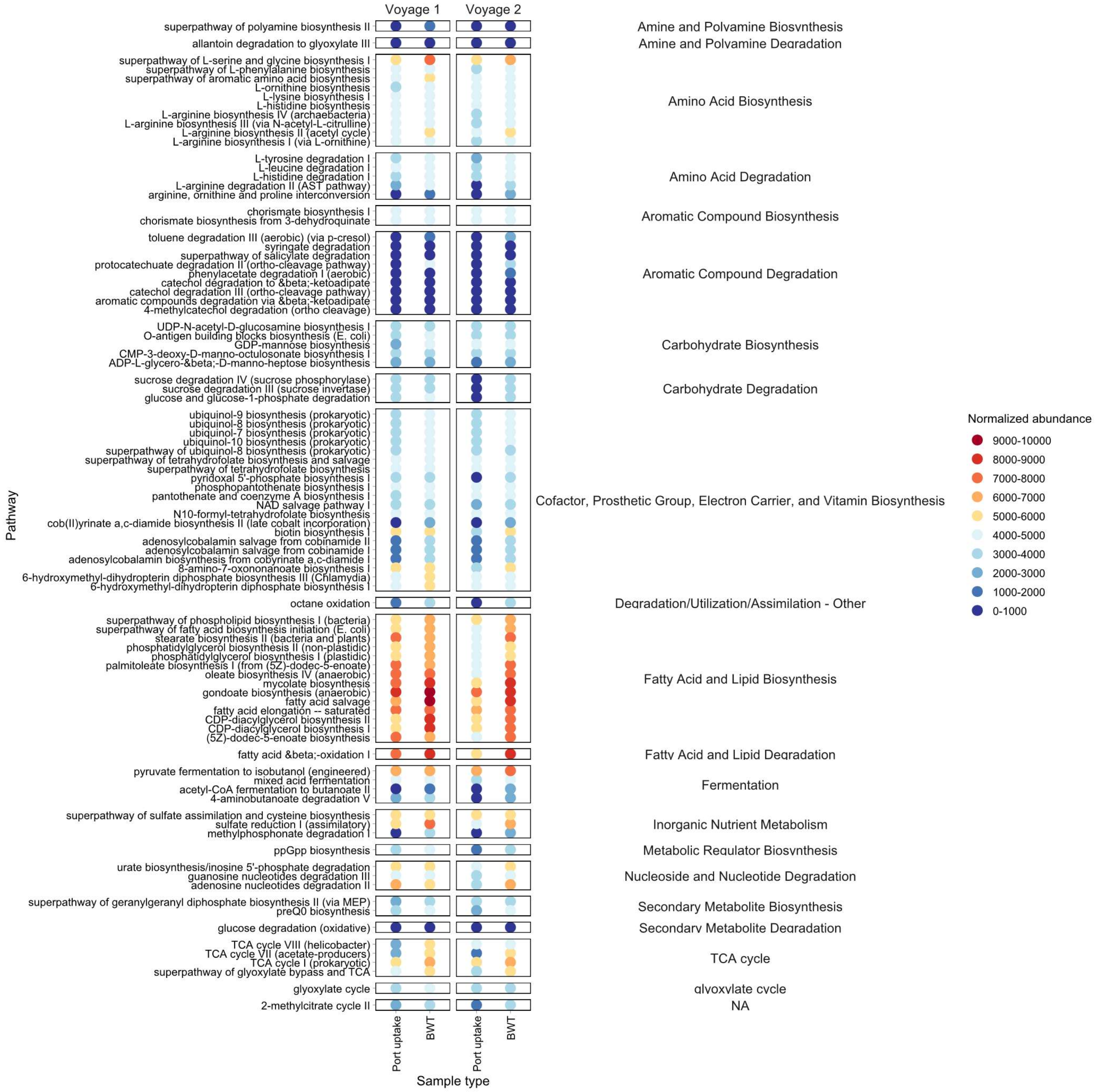


Figure S4. The normalized abundance of all inferred metabolic pathways that increased in abundance in BWT samples when compared to port uptake samples.
